# From lichens to crops: Pathogenic potential of *Pseudomonas syringae* from *Peltigera* lichens is similar to world-wide epidemic strains

**DOI:** 10.1101/2023.11.30.569175

**Authors:** Natalia Ramírez, Emma Caullireau, Margrét Auður Sigurbjörnsdóttir, Elodie Vandelle, Oddur Vilhelmsson, Cindy E. Morris

## Abstract

The presence of bacteria belonging to the *Pseudomonas syringae* complex (*P. syringae*) in the natural vegetation of several Icelandic habitat types has been recently reported, raising questions about the risk to Icelandic crops, particularly given the expected increase in agricultural activity due to climate warming. This study takes advantage of Iceland’s unique characteristics and the discovery of *P. syringae* in *Peltigera* lichens to gain a better understanding of the potential risk posed by this newly discovered ecological niche. The main objective is to evaluate the pathogenic potential and fitness in crops of *P. syringae* strains isolated from *Peltigera* lichen sampled in Iceland, focusing on strains that belong to phylogroups 1 and 2, which commonly contain epidemic strains. The results indicate that *P. syringae* isolated from Icelandic *Peltigera* lichen have a comparable fitness to epidemic strains in eight out of ten tested plant species. Furthermore, pathogenicity assessment on three plant species highlighted that certain strains also caused similar symptoms and disease severity compared to epidemic strains. These findings provide valuable insights into the potential risks posed by *P. syringae* from Icelandic natural habitats and illustrate how strains from these habitats have a wide pathogenic potential to crops without having encountered these crops in the last several thousand years of their presence in Iceland.

## Introduction

Bacterial strains within the *P. syringae* complex are present in connection with diverse biotic and non-living substrates worldwide. The ability of *P. syringae* to adapt to a wide range of habitats linked to the water cycle is thought to be a driver of its broad host range (Morris et al. 2013). This idea is reinforced by the observation that numerous strains from this group of bacteria isolated from non-agricultural environments are phylogenetically closely related to plant-associated strains and have also been shown to be pathogenic on plants such as kiwifruit and tomato (Morris et al, 2019). For this reason, recent research has continued to explore the ecology and pathogenicity of *P. syringae* outside the context of crops.

According to Ellis et al. (2010), agricultural land is becoming increasingly prevalent compared to other vegetated areas on Earth. However, Iceland stands out as an atypical region with a limited amount of cultivated land, thereby providing a unique opportunity to study the adaptation of *P. syringae* without a predominant influence of local agriculture. Iceland’s flora is characterized by a relatively small number of native species of vascular plants, comprising around 530 species (Wąsowicz 2020). Additionally, the country is home to a diverse range of lichens (755 species) (Kristinsson and Heiðmarsson, 2009) and mosses (around 600 species) (Jóhannsson 2003). However, the vegetation of Iceland evolved in the absence of large herbivores and subsequently is vulnerable to grazing and human activities (Runólfsson 1987). Furthermore, Iceland’s position on the border between Arctic and Atlantic waters and air masses, known as the polar front, makes for an interface (Jónsdóttir et al. 2005) which creates favourable climate conditions for *P. syringae* with average temperatures between 2 to 14°C throughout the year (Ogilvie and Jónsson, 2001).

A lichen is a composite organism resulting from a symbiotic association between a fungus, known as the mycobiont, and one or more photosynthetic partners, termed photobionts. The photosynthetic partners are typically green algae or cyanobacteria. This mutualistic relationship forms a unique structure known as a thallus, which is the visible body of the lichen. (Boustie and Grube, 2005; Feuerer and Hawksworth, 2007; Cardinale et al, 2006; Leiva et al, 2021), fungi, sometimes pathogenic for the lichen (Bates et al, 2012; Spribille et al, 2016), archaea (Bjelland et al, 2011; Garg et al, 2016), and viruses (Eymann et al, 2017). While lichens heavily depend on the atmosphere for water intake, their remarkable resilience in challenging environments is partly attributed to the microbiome associated with lichens, playing a crucial role in their survival. (Leiva et al, 2021; Pisani et al, 2011; Bates et al, 2011; Bjelland et al, 2011; Cardinale et al, 2012, 2008, 2006; Grube and Berg, 2009; Hodkinson and Lutzoni, 2010; Mushegian et al, 2011; Selbmann et al, 2010; Sigurbjörnsdóttir et al, 2015).

Previous research has shown that *P. syringae* is prevalent in Iceland on wild vascular plants and moss (Morris et al. 2022), confirmed by the observations of *P. syringae* genes in the lichen metagenome of *Peltigera membranaceae* (Sigurbjörnsdóttir et al. 2015; Sigurbjörnsdóttir, 2016). This prompted researchers to investigate its ubiquity across several types of plants and lichens in Iceland, hypothesizing that lichens may serve as non-host reservoirs for *P. syringae* (Vilhelmsson et al. 2016). Morris and colleagues (Morris et al. 2022) unveiled how the genetic lines of *P. syringae* in the Icelandic region are monophyletic, indicating that they may have evolved separately from the *P. syringae* populations elsewhere in the world during the relatively short geological history of Iceland. However, these monophyletic haplotypes represent different phylogroups (PGs) (Morris et al. 2022). This illustrates the extraordinary adaptive properties throughout the *P. syringae* complex.

*P. syringae* was found in lichens in Iceland, specifically in species of the genus *Peltigera* (Ramírez et al. 2023). To delve deeper into this discovery, a phylogenetic analysis of *P. syringae* strains collected from various sources, including lichens, tracheophytes, and moss was conducted. The analyses revealed significant differences among strains between geographical locations, showing a greater similarity of *P. syringae* within a site across all vegetation types rather than within vegetation types across sites. Moreover, *Peltigera* thalli harbored a consistent population density of *P. syringae* although it was lower than that on moss and tracheophyte samples (Ramírez et al. 2023). This finding underscores the adaptability of *P. syringae* to inhabit a diverse range of vegetation beyond higher plants, offering novel insights into its evolutionary dynamics.

Assigning *P. syringae* strains to phylogroups based on their citrate synthase gene sequences (Berge et al. 2014) is a useful tool for understanding their phenotypic variations. Many of the strains isolated from *Peltigera* lichens can be assigned to phylogroups PG01 and PG02. These phylogroups contain a wide range of epidemic strains, as documented by Berge and colleagues (Berge et al. 2014). Roughly fifty percent of the lichen thalli were found to host PG01 and/or PG02 strains. Among the *P. syringae* isolates, PG02 included approximately 14% of strains and PG01 accounted for a mere 4% of the overall *P. syringae* population derived from *Peltigera*, while the remaining 82% were assigned to the environmental habitat-associated phylogroups PG10 and PG13 (Ramírez et al. 2023).

In light of the ubiquity of bacteria in the *P. syringae* complex on vegetation in Iceland and, in particular, the presence of the PG01 and PG02 phylogroups showing high frequencies of strains displaying a functional T3SS (Berge et al. 2014), our goal was to assess the fitness and pathogenic potential – mainly on crops - of strains from PG01 and PG02 from Icelandic *Peltigera* compared to strains in the same phylogroup isolated from epidemics on crops elsewhere in the world.

## Materials and methods

### Bacterial strains

*P. syringae* strains belonging to PG01 and PG02 were randomly selected from a collection of strains isolated from *Peltigera* lichen that previously tested positive in a HR test to represent the range of diversity of these phylogroups associated with this group of lichens (Fig. 1). As positive controls, strains belonging to PG01 and PG02 from epidemic occurrences were chosen due to their well-established pathogenic potential and consistent behavior, as revealed by previous work (Morris et al. 2019). For fitness assessment, the reference strains included CC0094 (PG02d) and Pto DC3000 (PG01a). For the evaluation of pathogenicity, CC0125 and CFBP1906 (PG02b), and CC0094 (PG02d), were employed as reference controls, aligning with the findings outlined previously (Morris et al. 2000).

**Fig. 1.**
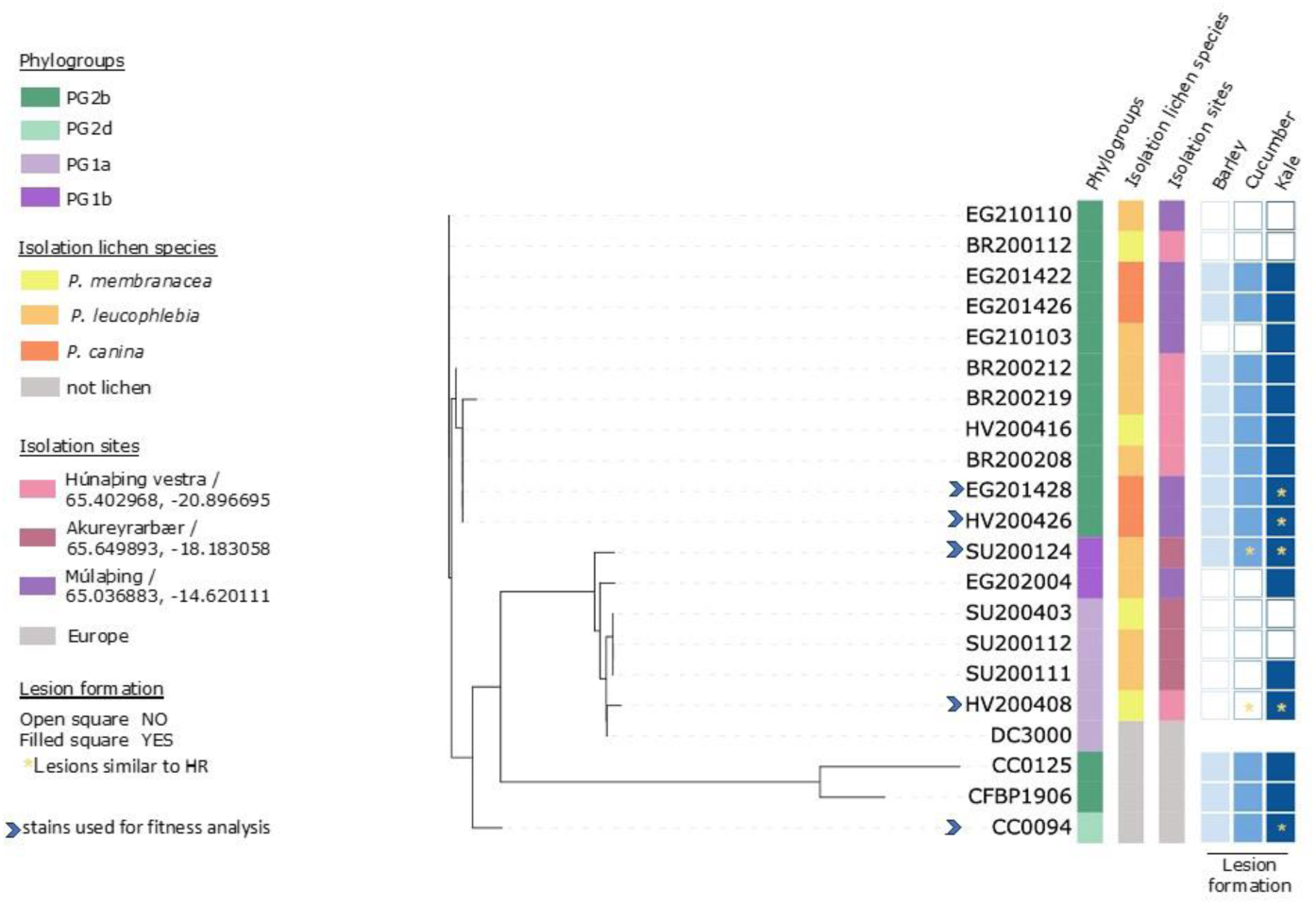
Phylogenetic tree illustrating the strains included in the study, along with details on lichen of isolation, phylogroup classification, site of isolation, and lesion formation in kale, cucumber, and barley. The color code represents different categories, and an arrow indicates the strains selected for fitness analyses.

### Inoculum preparation

Bacterial inoculum was prepared from 24-72 hr growth on King’s B (KB) medium (King et al. 1954). A loopful of growth was resuspended in phosphate buffer (TP1: 8,75g of K_2_HPO_4_ and 6,75g of KH_2_PO_4_ diluted in 1L of distilled water) and adjusted with a spectrophotometer to 10^8^ CFU mL^−1^ (OD_600 nm_ = 0.1). The inoculum was further diluted in phosphate buffer, resulting in a concentration of 10^7^ CFU mL^−1^ for the pathogenicity tests and 10^6^ CFU mL^−1^ for the in planta in the fitness tests. Inoculum concentration was verified by dilution plating.

### Plant material

For fitness tests, plant species belonging to 10 families were used: rice (*Oryza sativa*), tomato (*Solanum lycopersicum*), thale cress (*Arabidopsis thaliana*), annual mugwort (*Artemisia annua*), spinach (*Spinacia oleracea*), garlic chives (*Allium tuberosum*), tobacco (*Nicotiana tabacum*), kale (*Brassica oleracea*), cucumber (*Cucumis sativus*) and barley (*Hordeum vulgare*). Pathogenic potential was evaluated on the last three plant species, which are cultivated in Iceland. Cultivar details, growing conditions, and cultivation dates can be found in Table S2. The duration of cultivation in the greenhouse was tailored to the specific plant species. In the greenhouse environment of Montfavet, France, the plants were cultivated during a period spanning from September 2022 to January 2023. Adequate watering was provided in accordance with the plants’ needs. Plants were grown in TS3 substrate mixture from Klasmann-Deilmann (Geeste, Germany).

### Evaluation of fitness *in planta*

Fitness of strains in plants was determined in terms of population growth after inoculation using a modified protocol based on methodologies outlined in Clarke et al,(2010) Donati et al,(2020) and Kim et al, (2022). Leaf tissue was wounded and strains were inoculated into plants by placing a 5µL-drop of inoculum (10^6^ CFU mL^-1^), letting the inoculum be absorbed by the plant. An emery board/emery paper was used gently to create a fresh wound on the surface of the leaf. Each plant received a single inoculation, and there were 10 replicate plants per strain.

Three to five plants were inoculated with phosphate buffer as negative control for each plant species. For a given plant species, all strains were inoculated simultaneously to facilitate between-strain comparisons. Via a randomized complete block design, the plants were set up in a growth chamber and incubated at 24 °C (14 h light, 10 h dark) and 75% humidity.

At 1 and 8 days post inoculation (dpi), five leaves were collected from each plant species for each strain, except in the case of *A. thaliana*, where six leaves were collected. Prior to maceration, leaves bearing soil residues were gently cleaned using dry paper. Each leaf was stomached for 2–3 min in sterile phosphate buffer (10µL of 0.1 M buffer per mg of fresh plant weight amounting to between 1-30 ml depending on the leaf size) in sterile stomacher bags. Aliquots of serial dilutions of the macerate were plated on KB medium. The plates were placed in a dark environment and incubated at room temperature for a period of 48-72 hours. *P. syringae* colonies were identified based on their shape, size, color, texture, elevation, and margin (Morris et al. 2022). The abundance of *P. syringae* in the inoculated leaves was expressed per leaf.

### Inoculation, incubation, and disease assessment for pathogenicity assays

We employ a modified protocol inspired by Bartoli et al,(2015) and Cazorla et al, (1998). One leaf per plant was infiltrated with a needleless syringe with 5-10 μL of inoculum (10^7^ CFU mL^−1^). Six to eight replicate plants were utilized per plant species per strain. The negative control consisted of phosphate buffer. Plants were arranged in a growth chamber in a randomized complete block design and incubated at 24 °C (14 h light, 10 h dark) and 75% humidity for 14 days.

Symptoms were scored at 2, 5, 9 and 14 dpi in terms of the maximum length of the necrosis that developed at the inoculated site except for cucumber that was observed only until 9 dpi. Symptoms were also photographed and described at each scoring date. Strains were classified as pathogenic if the necrotic length was significantly different than the negative control.

To verify that symptoms were caused by *P. syringae*, isolations were made from at least three infected tissue replicates per strain. Sections of tissue (at the interface of necrotic and the green surrounding tissue) were aseptically collected and placed on KB plates and incubated at 24°C in darkness. After 2-3 days, colony morphology, colour and fluorescence under 366 nm were assessed.

### Statistical analyses

Utilizing Microsoft Excel and R Studio, statistical analyses were conducted. This included t-tests, graph plotting with the ggplot2 package, and one-way ANOVA. The t-test was applied to determine the differences between strains inoculated into the same plant species at various time points. Moreover, one-way ANOVA was employed, followed by post hoc Tukey-Kramer analysis to unravel strain variations.

## Results

### *P. syringae* isolated from Icelandic *Peltigera* lichens and epidemic strains have similar fitness in plants

The fitness analysis revealed that *P. syringae* strains isolated from Icelandic *Peltigera* lichen (hereafter referred as lichen strains) exhibited overall population growth levels similar to the epidemic strains CC0094 and DC3000 at both 1 and 8 dpi under the specified culture conditions across the ten plant species tested, except for cucumber, barley and tomato. Indeed, differences emerged in tomato at 1 dpi, with the population size of *Peltigera* lichen *P. syringae* strains reaching higher densities compared to epidemic ones (Figure S1), although no difference was further observed at 8 dpi. Conversely, despite a similar population density at 1 dpi, the *Peltigera* strains showed a lower population level at 8 dpi compared with epidemic strains in cucumber and barley (Fig. 2). For the latter, however, considering both time points, the comparison of the total population of lichen strains was significantly higher than the population density of epidemic strains (p value < 0.05). This indicates that *Peltigera* strains display, as whole, the same *in-planta* fitness than epidemic strains, with some variability among plant species.

**Fig. 2.**
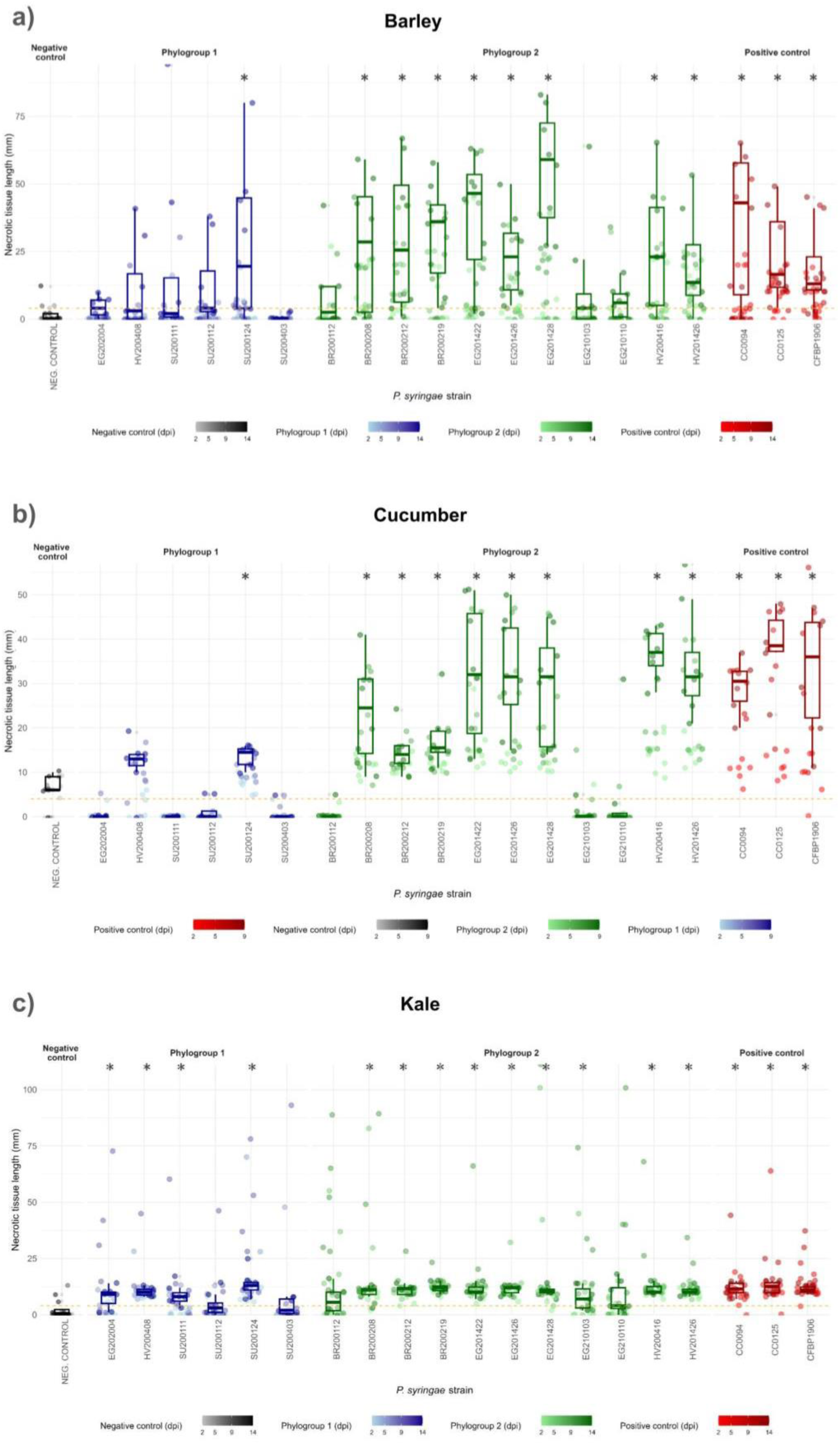
Necrotic tissue length in mm on leaves infiltrated with *P. syringae*. Dots represent all measurements at different time points distinguished by the darkness of the color and the green color correspond to PG1 strains and PG2 was colored in blue while the boxplots summarize the necrotic length on: a) Barley at 14 dpi; b) Cucumber on 9 dpi and; c) Kale on 14 dpi. All plants were incubated at 24 °C (14 h light, 10 h dark) and 75% humidity for up to 14 days. The *P. syringae* strains that exhibit statistical significance from the negative control (*p* value < 0.05) are marked with an asterisk on this graph

Interestingly, each lichen strain behave like the epidemic one belonging to the same phylogroup. Indeed, the main differences were observed more at phylogroup level, with PG02 (including CC0094, HV201426 and EG201428) showing higher bacterial densities compared to PG01 (DC3000, SU200124 and HV200408), at 1 dpi only, in 7 out of the 10 analyzed plant species. Such discrepancy, that was not noticed anymore at 8 dpi, suggests a better capacity of PG02 strains to adapt and start growing in plants independently of their origin.

On the other hand, when examining strains individually, at 1 dpi, CC0094 consistently showed a greater population size compared with DC3000 in all plant species but *Arabidopsis*, and higher than all strains but HV200426 in tobacco (Fig. 2). Conversely, at 8 dpi, distinctions were observed only in cucumber, with DC3000 having greater population sizes than all lichen strains, and in barley, where CC0094 had a significantly higher population size than SU200124 (p-value <0.05) (Fig. 2). Finally, looking at the dynamic of population growth, we observed that, in general, all strains (epidemic and lichen) display a population increase between 1 dpi and 8 dpi in all plant species, except for HV200408 and SU200124 that decreased over time in cucumber (Fig.S2), while no strain but EG210103 did grow in kale between 1 and 8 dpi (Fig.S3).

### *P. syringae* from *Peltigera* and from crop epidemics have similar pathogenic potential

For this analysis the set of lichen strains was enlarged to a total of 17, including the 4 strains used for *in-planta* fitness investigation, together with 4 and 9 additional strains belonging to PG01 and PG02, respectively. To account for non-specific reactions of the plants due to wounding, the size of the necroses on plants inoculated with selected strains was compared to those observed on control plants. Nine out of the 17 strains of *P. syringae* isolated from lichen produced necrotic symptoms that were significantly longer than any of the necroses observed on the negative controls in barley (Fig. 3.a) and cucumber (Fig. 3.b), and comparable to the lesions caused by epidemic strains. These same nine *Peltigera* strains and four additional strains, for a total of 13 out of 17, also caused lesions that were significantly longer on kale when compared to damage on control plants, and in the same size range than those observed with epidemic strains (Fig. 3.c).

**Fig. 3.**
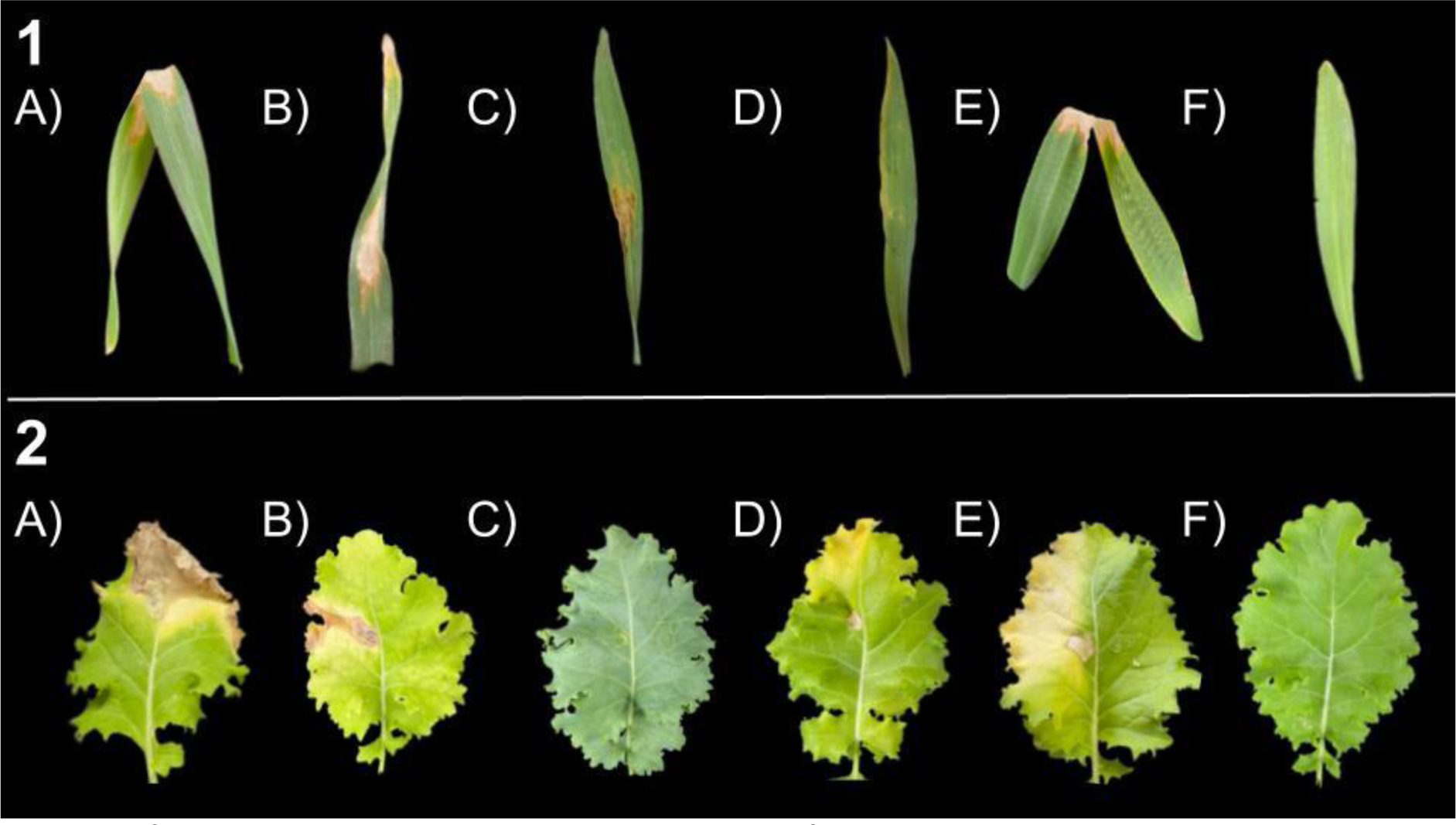
Leaf symptoms at 14 dpi. The leaves represent some of the most common symptoms observed on each plant species.1. Barley infected with strains a) HV201426; b) EG201422; c) EG201428; d) HV201426 e) Positive control CFBP1906; and f) Negative control. 2. Kale infected with strains a) HV201426; b) EG210110; c) SU200403; d) EG201426; e) Positive control CFBP1906; and f) Negative control

According to lesion size, symptoms were most severe on barley and cucumber compared to kale, although we cannot rule out the possibility that such differences could be attributable to technical aspects (e.g, plant growth conditions, plant age). Moreover, not only the number of PG02 strains (7 in barley and cucumber, 9 in kale) causing significant lesions was higher than the number of PG01 strains (1 in barley and cucumber, 4 in kale) but, considering only necrosis-inducing strains, symptoms caused by PG02 strains were overall more severe than those caused by PG01 strains. Indeed, symptom severity (lesion length) in most cases was within the same range as those caused by the strains from crop epidemics (Fig. 3).

In barley, in addition to necrosis, other symptoms such as chlorosis, leaf collapse, wilting, water-soaked areas or a slightly pale appearance were also recorded, mainly following infection with necrosis-inducing strains (Fig. 4.1). Similarly, the infiltration of the *Peltigera* strains in kale caused symptoms such as chlorosis, water-soaked areas sometimes surrounded by a yellow halo and pale appearance, besides necrotic lesions (Fig. 4.2). The observed symptoms in cucumber included necrosis, chlorosis, and lesions with a yellow halo (images not available).

**Fig. 4.**
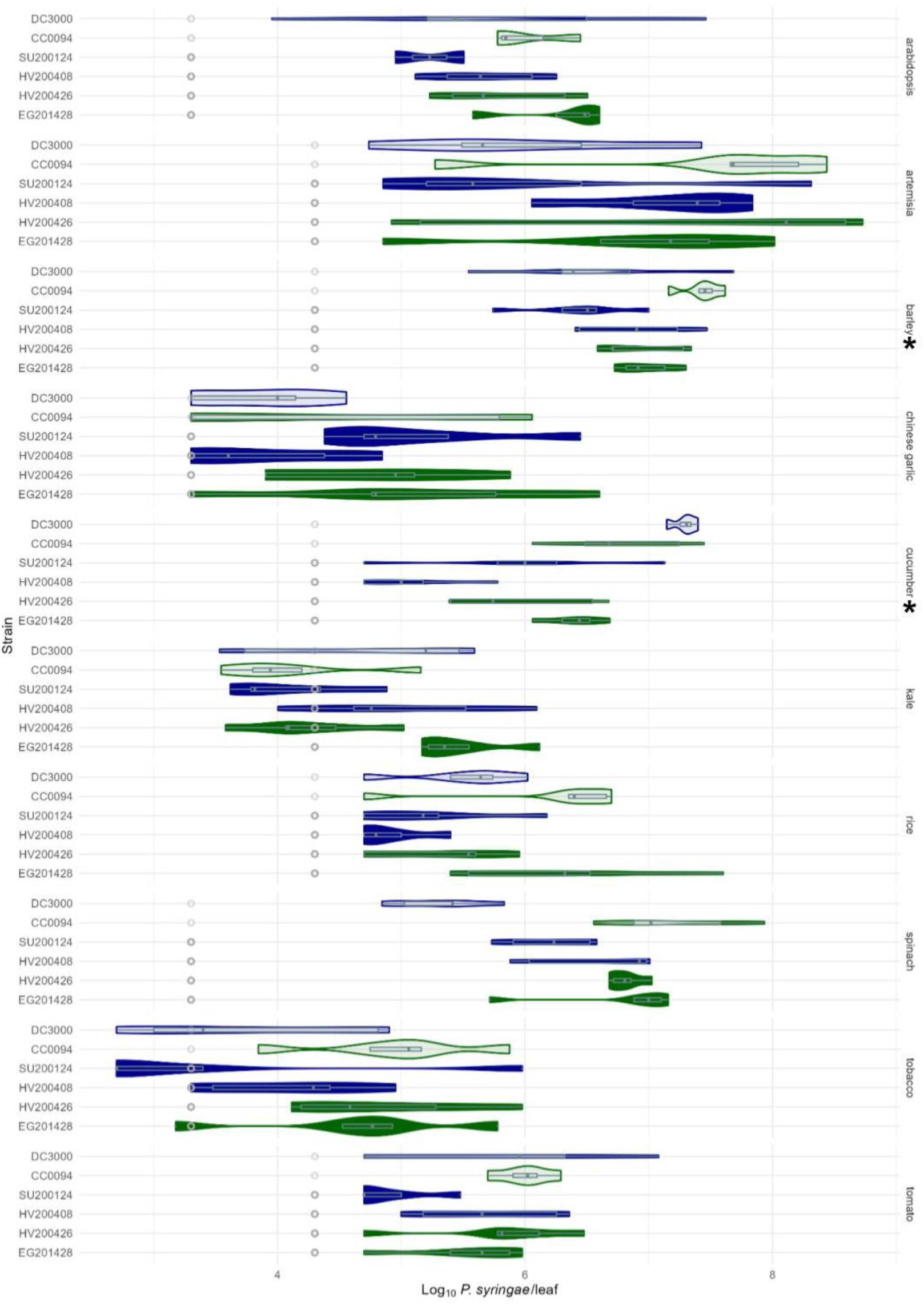
*P. syringae* population in inoculated leaves of ten plant species on the 8th day post-infection. Violin and boxplot graphs resume the values of *P. syringae* per leaf for 5 or 6 replicates per strain inoculated in each plant. The Icelandic strains isolated from lichens are represented by a solid color while the epidemic strains used as control are plotted as transparent. PG1 strains are represented by the color blue, while PG2 strains are depicted in green. Grey dots indicate the minimum detection level. Asterisk denotes a plant species where statistically significant differences in *P. syringae* population sizes between epidemic and *Peltigera* strains were observed

In some cases, however, necroses resembled hypersensitive-like cell death, *i.e.* occurring shortly after infiltration and apparently localized to the site of infection. In particular, such a phenotype was observed in cucumber inoculated with the *Peltigera* strains belonging to PG01, namely HV200408 and SU200124, which induced smaller lesions, reaching a maximum length of around 15 mm, compared to the lesions induced by PG02 strains with a lesion size comprised between 30 and 40 mm (Fig.S2). Interestingly, the two above-mentioned strains showed a decrease in population size in cucumber 8 dpi compared to 1 dpi, likely related to the induction of HR leading to the restriction of bacterial growth.

The induction of HR-like cell death was also common to numerous strains tested in kale, including the lichen strains HV200408 and SU200124 from PG01, and HV201426 and EG201408 belonging to PG02, together with the epidemic strain CC0094 (Fig.S3). Interestingly, the lesions induced by these strains were already clearly visible after only 2dpi, but did not increase further up to 14 dpi. Considering the absence of bacterial population increase between 1 and 8 dpi in fitness experiments (Fig.2), these results indicate the capacity of these *Peltigera* strains to induce a rapid HR in kale that an in turn stop bacterial growth. Conversely, the PG02 lichen strain EG210103 induced significant lesions in kale only after 5 dpi and these necroses were characterized by the presence of chlorosis (Fig.S3.b). Moreover, this strain was the only one capable of growing between 1 and 8 dpi in fitness experiments, thus supporting the absence of HR in this case.

## Discussion

Here we have revealed that *P. syringae* isolated from *Peltigera* lichen in Iceland has fitness and pathogenicity levels comparable to strains isolated from epidemic crops worldwide. As a component of pathogenicity, some *Peltigera* strains were capable of inducing HR. Hence, even though we do not know what the host range would be in a cropping system, the strains from lichens possess a functional system to deliver effectors and to be recognized by a plant as something more than a saprophyte.

Numerous studies have demonstrated that *P. syringae* retains its pathogenic capability even in the absence of significant agricultural pressures (Morris et al. 2008; Morris et al. 2013; Morris and Moury, 2019), especially in those strains that have a wider range of habitat. The hypothesis suggests that the pathogenicity of *P. syringae* could be more pronounced and evident in agricultural contexts due to the lack of genetic diversity of the host and some practices for cultivation that might favor a rapid growth of *P. syingae,* eventhough it can also grow in non-agricultural plants as *Arabidopsis thaliana*, grass and ornamental plants (Jones et al, 1986; Katagiri et al, 2002; Sato et al, 2001). However, the unique context of the Icelandic *P. syringae* strains that have evolved in Iceland for thousands of years makes them an interesting case study (Morris et al. 2022) as it illustrates the ancestral nature of the traits that confer pathogenicity and their maintenance in populations in natural habitats (Xin et al. 2018).

Although the pathogenicity observed in controlled conditions do not precisely reflect the pathogenicity of bacterial strains in the field, our results clearly illustrate the pathogenic potential of Icelandic strains, under favorable conditions. Thus, while this suggests the potential threat posed by *P. syringae* in Iceland, it is crucial to keep in mind that *P. syringae* is ubiquitous in the environment. In this light, it is notable that there have been no reported instances of *P. syringae* causing diseases in local crops in Iceland nor to the native vegetation. This highlights the role that environmental factors, such as cold temperatures that reduce the cultivation season or average temperatures under the optimal for *P. syringae* to develop aggressive traits (Bender et al. 1999), likely play in constraining bacterial proliferation and safeguarding plant health. The interplay of environmental conditions significantly contributes to the assessment of disease potential (Morris et al. 2023).

*P. syringae* in Iceland has evolved over the past 10000 years, at least, separate from contact with plants that are common to agricultural landscapes such as rice and tomato (Morris et al. 2022). Nevertheless, Icelandic strains display some level of fitness in these plants, which could be then considered as novel potential hosts for such stains. This raises intriguing questions about the establishment of host specificity. However, the aggressiveness detected for Icelandic *P. syringae* might be related to the adaptation to environmental habitats, which have been linked to the evolution of their pathogenicity (Morris et al. 2010).

As previously demonstrated for *P. syringae* overall (Morris et al, 2019), it has not been possible to make any inference of pathogenic capacity/host range based on the substrate of isolation (habitat or lichen species) or according to the phylogroup of the Icelandic strains (Fig.1). This is also consistent with the observation of the mixing of populations of *P. syringae* between crop and environmental habitats, whereby they are not genetically distinguishable as different populations (Monteil et al, 2016; Morris et al, 2010). Moreover, the lichen strains resemble more the strains of *P. syringae* at a same site than other strains from lichens at distal sites (Ramírez et al, 2023). Thus, overall, these observations demonstrate that the habitat from which a strain is isolated is not necessarily the habitat on which it has spent the most time and via which selection pressures might have dominated.

The finding of *P. syringae* raised concerns about a possible threat to Icelandic crops in case the strains isolated showed pathogenic properties. Icelandic cool climate can play an important role in pathogenicity of *P. syringae*, due to tissue damage that intensified by frost episodes and humid environments, as it has been shown that the presence of a layer of free water is essential for infection (Lamichhane et al. 2015). Furthermore, certain studies have indicated that lower temperatures enhance the functionality of T3SS (Type III Secretion System) and T6SS (Type VI Secretion System), which are typically associated with increased aggressiveness (Tribelli and López, 2022; Puttilli et al, 2022). Despite the limited agricultural land in Iceland (Denk et al. 2011), certain crops are cultivated on an industrial scale in greenhouses, with yields exceeding 1000 tones annually. These include tomatoes and cucumbers, while outdoor cultivation involves crops like barley and potatoes. Nevertheless, the anticipation is that the range of plant species grown outdoors and their yields in Iceland will rise over the coming decades due to the predicted rise in temperatures (Parry 1991). The expansion of cultivable land, coupled with the anticipated milder, yet still relatively cold temperatures in Iceland, may pose a potential threat to Icelandic crops.

The results of this study are one more example of *P. syringae* strains isolated from environmental habitats, where they are not causing any obvious damage, or where they might even be beneficial (Morris et al. 2013). However, these strains can affect some plant species under specific conditions – conditions that might be increasingly probable with the changing climate and land use in Iceland.

## Supporting information

Figure S1 P. syringae population in inoculated leaves of ten plant species on the 1st day post-infection. Violin and boxplot graphs resume the values

Figure S2. Necrotic tissue length in cucumber a) across all strains at 2 dpi.

Figure S2. Necrotic tissue length in cucumber b) in strains displaying HR lesions at 2, 5, and 9 dpi.

Figure S3. Necrotic tissue length in kale a) across all strains at 2 dpi.

Figure S3. Necrotic tissue length in kale b) in strains displaying HR lesions at 2, 5, 9, 14 dpi.

## Ethics and integrity policies

- The authors declare no conflict of interest.
- This work was funded by the Rannis Icelandic Research Fund under project number 1908-0151.

