## Supplementary figures and images for "From lichens to crops: Pathogenic potential of *Pseudomonas syringae* from *Peltigera* lichens is similar to world-wide epidemic strains"

### Figure S1 P. syringae population in inoculated leaves of ten plant species on the 1st day post-infection. Violin and boxplot graphs resume the values

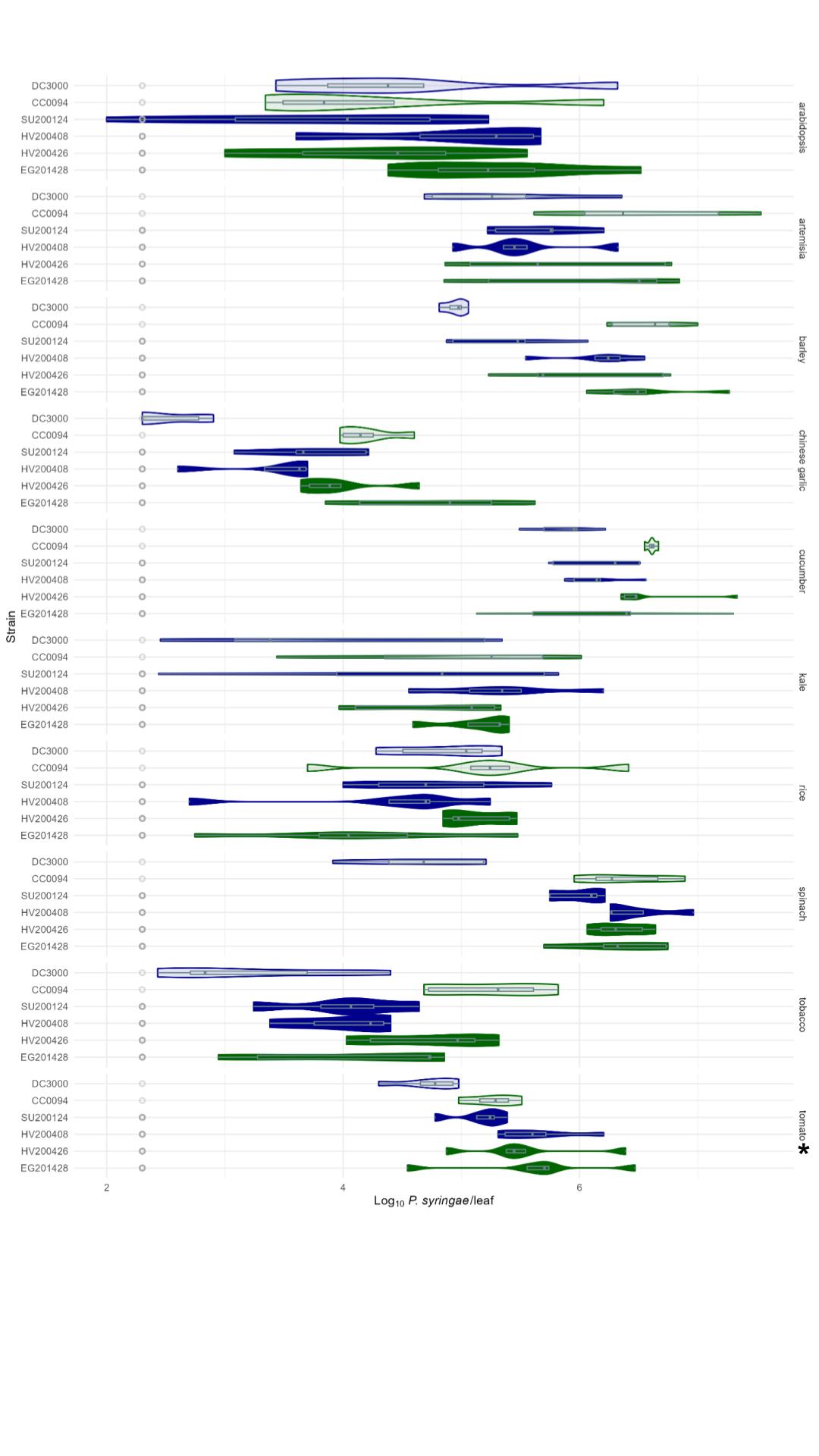

### Figure S2. Necrotic tissue length in cucumber a) across all strains at 2 dpi.

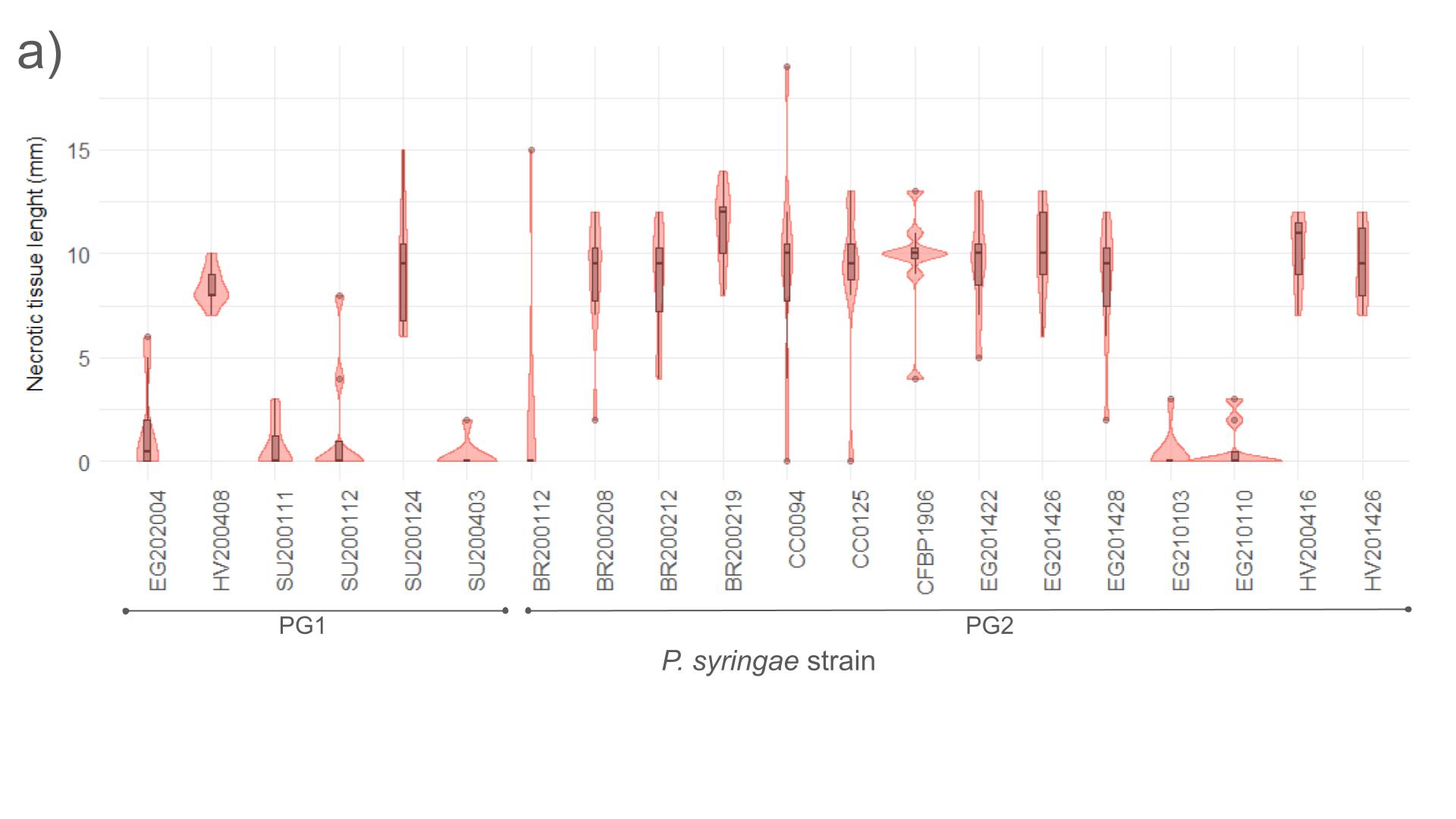

### Figure S2. Necrotic tissue length in cucumber b) in strains displaying HR lesions at 2, 5, and 9 dpi.

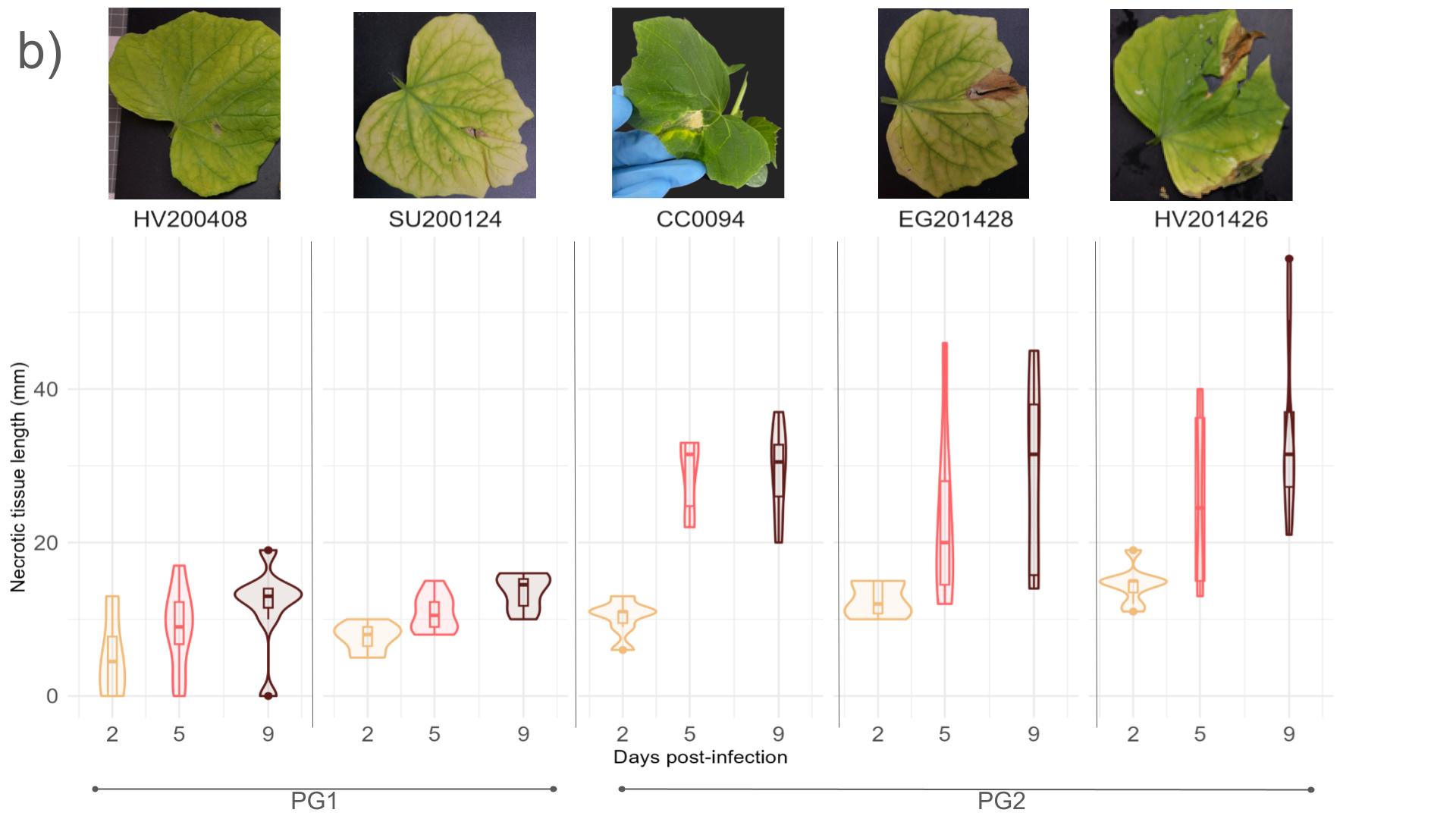

### Figure S3. Necrotic tissue length in kale a) across all strains at 2 dpi.

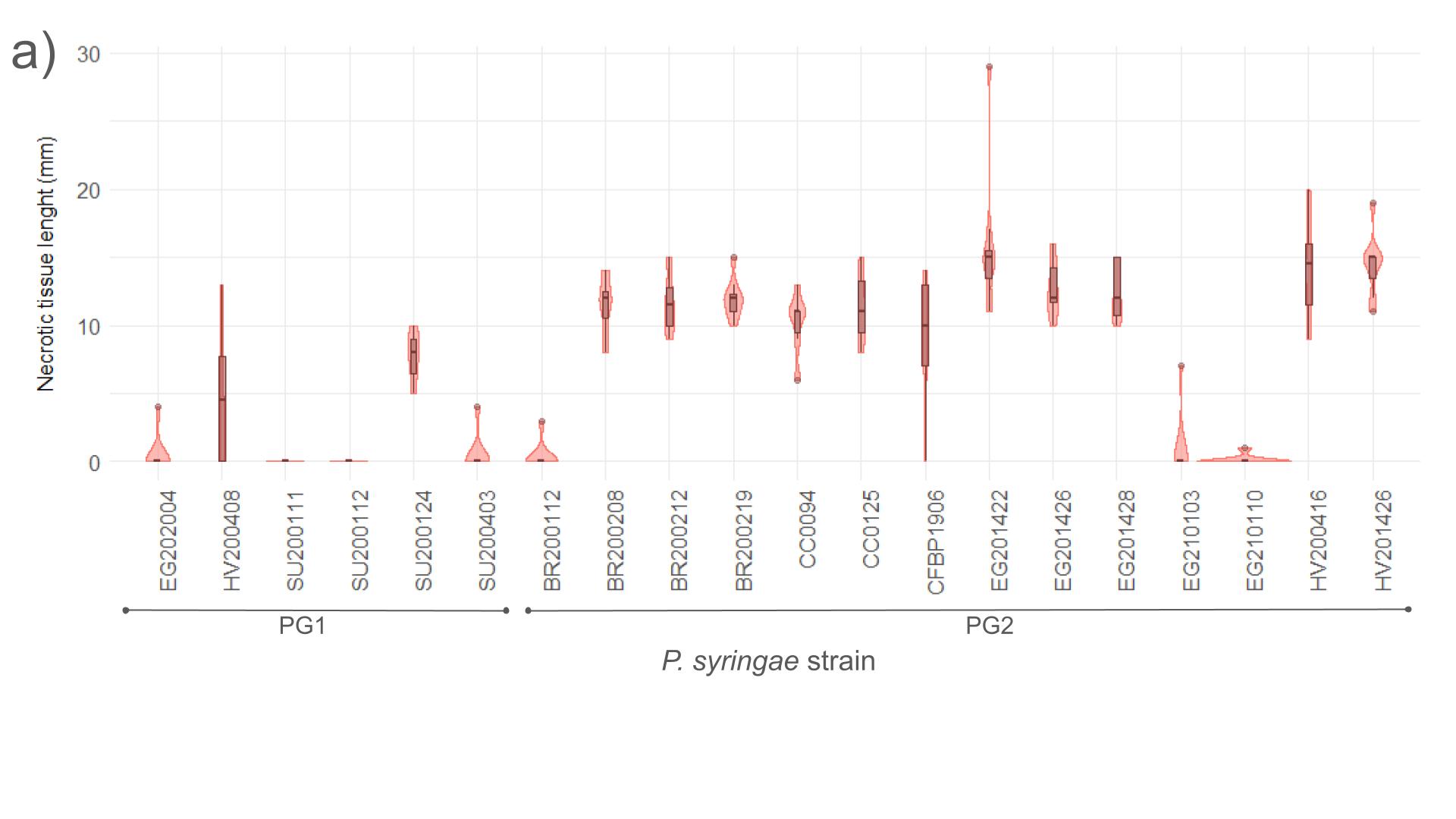

### Figure S3. Necrotic tissue length in kale b) in strains displaying HR lesions at 2, 5, 9, 14 dpi.

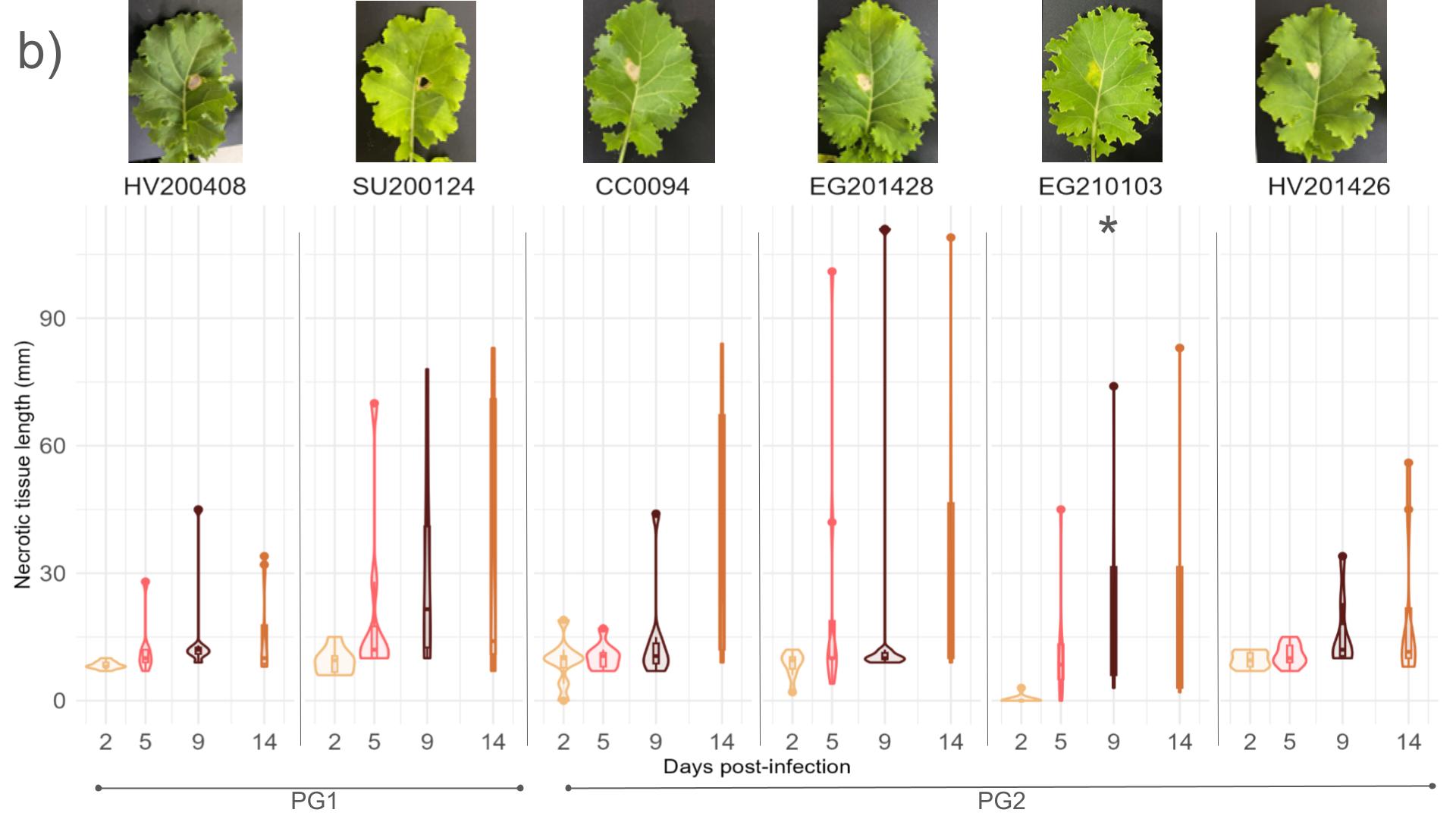
